# High-throughput functional annotation of influenza A virus genome at single-nucleotide resolution

**DOI:** 10.1101/005702

**Authors:** Nicholas C. Wu, Arthur P. Young, Laith Q. Al-Mawsawi, Anders C. Olson, Jun Feng, Hangfei Qi, Shu-Hwa Chen, I-Hsuan Lu, Chung-Yen Lin, Robert G. Chin, Harding H. Luan, Nguyen Nguyen, Stanley F. Nelson, Xinmin Li, Ting-Ting Wu, Ren Sun

## Abstract

A novel genome-wide genetics platform is presented in this study, which permits functional interrogation of all point mutations across a viral genome in parallel. Here we generated the first fitness profile of individual point mutations across the influenza virus genome. Critical residues on the viral genome were systematically identified, which provided a collection of subdomain data informative for structure-function studies and for effective rational drug and vaccine design. Our data was consistent with known, well-characterized structural features. In addition, we have achieved a validation rate of 68% for severely attenuated mutations and 94% for neutral mutations. The approach described in this study is applicable to other viral or microbial genomes where a means of genetic manipulation is available.

## Introduction

The influenza virus causes several hundred thousand deaths every year, and this number can reach millions in pandemic years. The huge socioeconomic associated with influenza highlights the importance of understanding of virus-host interactions [1, 2]. The rapidly evolving nature of influenza challenges the development of anti-influenza drugs and vaccine [3–7]. Consequently, it is important to develop drugs or vaccines that target indispensable regions on the influenza virus to maximize the genetic barrier for the emergence of resistance or escape mutations. Nevertheless, genetic research on the influenza virus has largely relied on naturally variants and individual mutants created in the laboratory. A substantial part of the genome remains uncharacterized.

Traditional genetics studies the relationship of a single genotype-phenotype at a time, and has been extensively to study panels of influenza mutations. However, the low throughput of traditional genetics limited the number of mutations being examined. In contrast, high-throughput genetics interrogates the phenotypic outcomes of multiple mutants in parallel. Genome-wide insertional mutagenesis is a common high-throughput genetics approach. It has been employed in the influenza virus to systematically identify regions that are tolerate to mutations [8]. However, the resolution of insertion-based approach is limited at the protein subdomain level. This resolution is insufficient to identify residues critical for replication. As a result, there is a demand for a high-throughput genetics platform at a single-residue resolution.

Recently, we have developed a high-throughput genetic platform which allowed us to profile the fitness effect of individual point mutations across the influenza A virus hemagglutinin (HA) segment [9]. The principle of the high-throughput genetic platform is to utilize a large mutant library and deep sequencing. Here, we extended this approach to quantify the fitness effects of each point mutation in 96% of the influenza A virus genome. This technique will enable systematic identification of indispensable regions for drug or vaccine targets. More importantly, it can be applied to any specified growth conditions for any virus that can be genetically manipulated.

## Results

### Quantification of the fitness effect of individual point mutation

Our high-throughput genetics platform aims to randomly mutagenize each nucleotide of the genome, monitor the changes in occurrence frequency for individual point mutations under specified growth conditions using massive deep-sequencing [9]. The changes in occurrence frequency of each point mutation (such as diminishment or enrichment) allow us to quantify the mutational fitness outcomes under the given growth conditions. The mutant libraries were created by error-prone PCR on the eight-plasmid reverse genetics system influenza A/WSN/1933 (H1N1) [10] (see materials and methods). Subsequently, eight viral mutant libraries were generated by transfection, each with one of the eight segments mutagenized. All viral mutant libraries were passaged for two 24-hour rounds in A549 cells (human lung epithelial carcinoma cells). The plasmid library and the passaged viral library were each sequenced by Illumina HiSeq 2000. Here, a relative fitness index (RF index) is used to estimate the mutational fitness effect. The RF index is calculated as:

RF index = occurrence frequency in passaged library)/(occurrence frequency in plasmid library)

The occurrence frequency of individual mutations was expected to be lower than the sequencing error rate (∼0.1%-1%) in next generation sequencing (NGS). Therefore, we utilized a two-step PCR approach for sequencing library preparation to distinguish true mutations from sequencing errors. In the first PCR, a unique tag was assigned to individual molecules. The second PCR generated multiple identical copies for individual tagged molecules. The input copy number for the second PCR was well-controlled such that individual tagged molecules would be sequenced ∼10 times. True mutations would exist in all sequencing reads sharing the same tag, whereas sequencing errors would not. Individual molecules, each carrying a unique tag, have an average copy number of ∼10 in the sequencing data, which validated the sequencing library preparation design (Fig. S1).

### Point mutation fitness profiling of influenza A virus genome

The RF indices for individual point mutations were profiled across 96% of nucleotide positions in the influenza A/WSN/1933 virus genome (Fig. 1). The remaining 4% of nucleotide were from the termini of each gene segment due to PCR amplification difficulty. As expected, a positive correlation exists between RF index and the degree of amino acid conservation of missense mutations (Fig. S2). In addition, the fitness data for well-characterized mutants were consistent with their phenotypes reported in the literature. Examples include a critical salt bridge for viral replication on nucleoprotein (NP) [11] (Fig. S3A), replication enhancement mutation on polymerase subunit (PB2) [12] (Fig. S3B), attenuation of oseltamivir resistance mutation on neuraminidase (NA) [13] (Fig. S3C), low fitness cost of amantadine/rimantadine resistance mutations on ion channel (M2) [5,14,15] (Fig. S3D), and the basic stretch on matrix protein (M1) required for assembly [16] (Fig. S4). Furthermore, comparison between our fitness data with the polymerase activity on 19 PB1 mutants previously reported showed an 80% correlation [17]. Mutants that displayed a severely attenuated (RF index <0.05) or neutral (RF index >0.4) phenotype were randomly selected across the genome, individually constructed and tested. The replication phenotype of each single mutant validated the profiling data with a confirmation rate of 68% for severely attenuated mutations and 94% for neutral mutations (Fig. 2). These data taken together provides validity to our fitness profiling data set.

**Figure 1.**
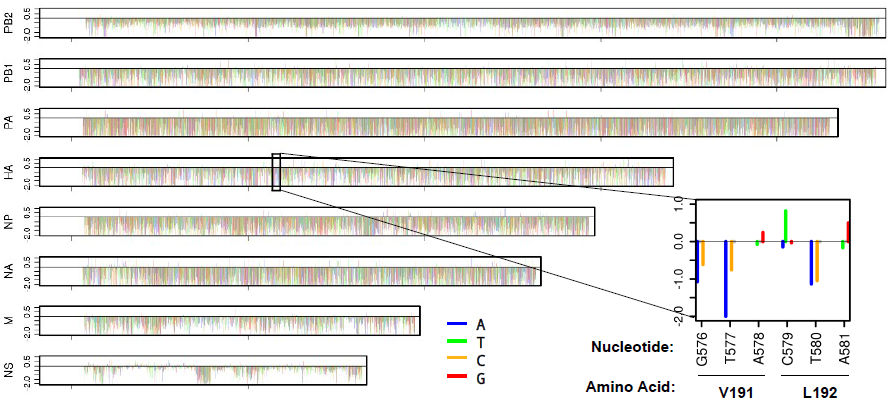
Single-nucleotide resolution fitness profiling. The RF index for individual point mutations across the genome was computed. Natural log of RF index, which is the ratio of occurrence frequency in the passaged library to the occurrence frequency in the plasmid library, represents the y-axis. Each nucleotide position is represented by four consecutive lines for the RF index that correspond to mutating to A (blue), T (green), C (orange), or G (red). The RF index of WT nucleotides is set as zero. Only point mutations with a coverage of ⩾ 30 tag-conflated reads in the plasmid library are shown. Point mutations with < 30 tag-conflated reads in the plasmid library is plotted as a gray dot on the zero baseline. The data track for HA is adapted from Wu et al. [9].

**Figure 2.**
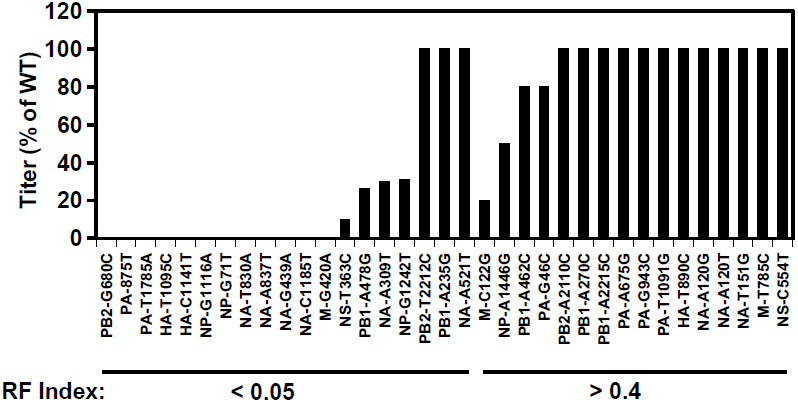
Experimental validation of severely attenuated and neutral mutations. Based on the data in Fig. 1, mutations that displayed a RF index of < 0.05 were classified as severely attenuated and > 0.4 were classified as neutral. Individual mutants were constructed and compared to the wild type (WT) replication phenotype. Post-transfection titers were plotted for lethal and viable mutants. Infection was initiated at an MOI of 0.05. Virus was harvested at 24 hours post infection. For the validated mutations with a RF index < 0.05, 68% have at least 1 log decrease in titer compared to WT. For the validated mutations with a RF index > 0.4, 94% have a titer within a 2-fold change as compared to WT. Overall the validation rate is ∼80%.

### Structural analysis and identification of indispensable protein surface

Our high-throughput profiling technique provides a basis to identify essential protein surfaces for drug targeting and indispensable regions for vaccine epitopes. We have performed a structural analysis on NA, a major influenza vaccine antigen. Here we identified a cluster of essential residues at the tetramer formation interface, suggesting that it bears functional importance and can possibly be a drug targeting site. In contrast, such a large cluster of essential residues could not be found in any other part of the NA surface. The lack of essential residues on the NA surface explain the functional basis of antigenic drift.

We have also performed a structural analysis using the PA subunit of the influenza virus RNA polymerase as an example to search for indispensable regions to aid in rational drug design. Increasing evidence suggests PA is a valuable target for drug development due to its polyfunctionality [18–20]. Our fitness data provided an informative reference for rational drug design. It captured several critical interactions between PA and PB1, such as the hydrogen bond between PA E617 and PB1 K11 (Fig. 3A), and the hydrophobic interaction between PA and PB1 via the volume-filling residues L666 and F710 (Fig. 3B). It has also revealed a cluster of essential residues on the PA surface consisting of eight amino acids (Fig. 3C), including K539 and K574, which were previously shown to be part of a lead compound binding pocket [19]. This patch of amino acids may be involved in an essential protein-protein interaction for viral replication. Similar analyses using our dataset have been applied to PA endonuclease domain and the M2 ion channel, which are plausible targets in drug development (Fig. S5-6). By projecting the fitness profiling data on three dimensional protein structures, it enables the identification of novel putative essential structural motifs that are surface exposed but not necessarily sequential in the primary sequence. This type of analysis reveals biological targets useful for rational drug and vaccine design. We propose that future antiviral drug design can incorporate the technique described in this study with in silico drug screening to increase the efficiency of therapeutic identification.

**Figure 3.**
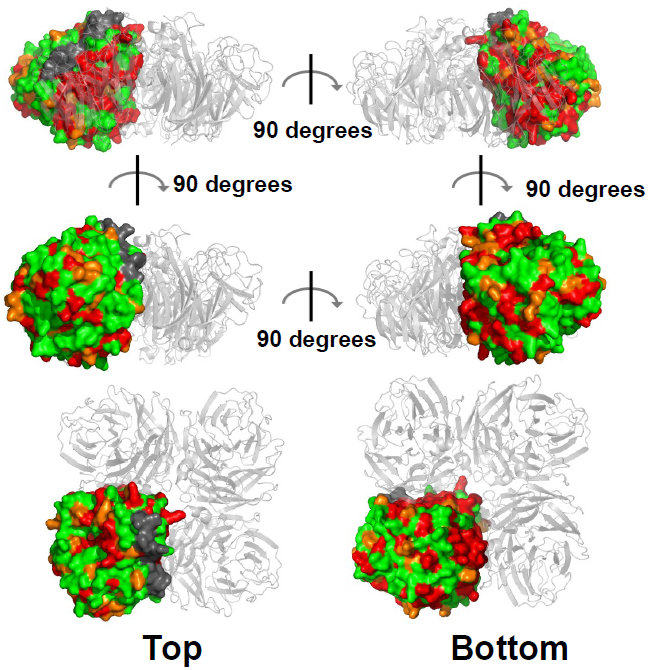
Structural analysis of the NA homotetramer interface. The RF index of the least destructive missense mutations for individual amino acids on the NA segment were projected on the protein structure (PDB: 3CL0) to identify for essential regions [28]. The RF index is color coded: RF index < 0.1, red; 0.1 ⩽ RF index < 0.2, orange; uncovered, grey. Only one monomer of the homotetramer is color coded.

**Figure 4.**
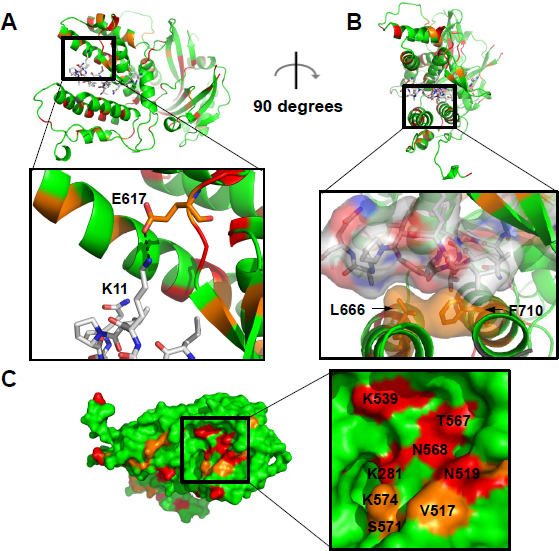
Structural analysis of the RNA polymerase PA subunit. The RF index of the least destructive missense mutations in the profiling data for individual amino acids on the PA segment are projected on the PA-PB1 complex crystal structure (PDB: 2ZNL) [29]. Most deleterious 10%, red; 10% to 20%, orange; Others, green. Our fitness data is capable to identify several critical interactions and putative functional sites. (D) A hydrogen bond between PA E617 and PB1 K11 is shown. Substitution of PA E617 is deleterious in our fitness data. (E) A hydrophobic interaction is shown between PA L666 and F710 and PB1. Substitution of L666 is deleterious in our fitness data. (F) A cluster of eight essential residues on the surface of PA is shown.

## Discussion

Sequence conservation was often taken as the sole parameter for identifying residues essential for viral replication, although conservation is not equivalent to essentialness for viral replication. It has been suggested that a significant fraction of conserved residues that are conserved in the influenza A virus are dispensable in viral replication [17,21,22]. In addition, new mutations were observed in every flu season, implying that residues that are naturally conserved currently may still be able to mutate under future unforeseen selection pressures. Therefore, a high-throughput fitness profiling complements the shortcoming in the sequence conservation analysis and allows identification of amino acid residues that are critical for viral replication in a defined cellular environment.

Here we provided a proof-of-concept study to profile the entire influenza A virus genome at single-nucleotide resolution. The fitness effects of individual point mutations were interrogated in a high-throughput manner by coupling a large mutant libary with NGS. However, the quantifiability of our platform can be further improved as sequencing technology advance. Similar experiments should be performed with strains across subtypes to identify mutations that display a genetic background-dependent fitness effect. These results would provide valuable information to dissect the evolutionary process of the influenza A virus. In addition, this platform can be applied to study the virus-host interaction under different cellular responses (such as apoptosis, autophagy, inflammasome induction, ER stress, etc.) and immune responses (such as NK cells, T cells, antibodies, macrophages, cytokines, etc.) that influence the viral replication in nature [23, 24]. Such results will significantly improve our understanding of the biological role of each residue on the genome of the influenza A virus. They will also help improve the design of a live attenuated influenza vaccine by minimizing the virulence. More importantly, it can potentially be adapted to other virus and microbes that can be genetically manipulated in the laboratory.

## Acknowledgments

We would like to thank J. Zhou, J. Yoshizawa, T. Toy and Z. Chen for performing the high-throughput sequencing experiment, K. Squire for support on data analysis, Y. Liu for advice on the PA structural analysis, Y. Liang and J. Bloom for valuable discussions.

## Materials and Methods

### Viral mutant library and point mutations

The plasmid mutant libraries were created by performing error-prone PCR on the eight-plasmid reverse genetics system of influenza A/WSN/1933 (H1N1) [10]. We PCR-amplified the flu insert with error-prone polymerase Mutazyme II (Stratagene, La Jolla, CA). Mutation rate of the error-prone PCR was optimized by adjusting the input template amount to avoid the accumulation of deleterious mutations. The restriction enzyme sites BsmBI and/or BsaI were added to the PCR primers, and used to clone into a BsmBI-digested parental vector pHW2000. Ligations were carried out with high concentration T4 ligase (Invitrogen, Grand Island, NY). Transformations were carried out with electrocompetent MegaX DH10B T1R cells (Invitrogen), and > 100,000 colonies for each segment library were scraped and directly processed for plasmid DNA purification (Qiagen Sciences, Germantown, MD). As extensive trans-complementation was expected during the transfection step, > 35 million cells were used for transfection to average out any bias or artifact generated from possible trans-complementation. Point mutants for the validation experiment were constructed using the QuikChange XL Mutagenesis kit (Stratagene) according to the manufacturer’s instructions.

### Transfections, infections, and titering

C227 cells, a dominant negative IRF-3 stably expressing cell line derived from human embryonic kidney (293T) cells, were transfected with Lipofectamine 2000 (Invitrogen) using 7 wildtype plasmids plus 1 mutant (library) plasmid. Supernatant was replaced with fresh cell growth medium at 24 hrs and 48 hrs post-transfection. At 72 hrs post-transfection, supernatant containing infectious virus was harvested, filtered through a 0.45 um MCE filter, and stored at −80°C. The TCID50 was measured on A549 cells (human lung carcinoma cells).

Virus from C227 transfection was used to infect A549 at an MOI of 0.05. Infected cells were washed three times with PBS followed by the addition of fresh cell growth medium at 2 hrs post-infection. Virus was harvested at 24 hrs post-infection. For the mutant library profiling, all viral mutant libraries were passaged for two 24-hour rounds in A549 cells. Our pilot experiments as well as our previous study revealed that two rounds of passaging were suffcient for profiling [25].

### Sequencing library preparation

DNA from the plasmid library or cDNA from the passaged viral mutant library were amplified with both forward and reverse primers each flanked with a 6 “N” tag and the flow cell adapter region. Flanking region for 5’ primer: 5’-CTACACGACGCTCTTCCGATCTNNNNNN-3’, Flanking region for 3’ primer: 5’-TGCTGAACCGCTCTTCCGATCTNNNNNN-3’. Following PCR, 93 amplicon products were pooled together. 15 million copies of the pooled product were used as the input for the second PCR, which was equivalent to 10 paired-end reads per molecule if 150 million paired-end reads (approximately one lane on an Illumina HiSeq 2000 machine) were sequenced. 5’-AATGATACGGCGACCACCGAGATCTACACTCTTTCCCTACACGACGCTCTTCCG-3’ and 5’-CAAGCAGAAGACGGCATACGAGATCGGTCTCGGCATTCCTGCTGAACCGCTCTTCCG-3’ were used as the primers for the second PCR. Products from the second PCR were submitted for NGS. The error-correction technique described in this study adapted the philosophy described for detecting rare mutations in human cells [26]. Raw sequencing data have been submitted to the NIH Short Read Archive under accession number: SRR1042008 (plasmid mutant library) and SRR1042006 (passaged mutant library).

### Data Analysis

Sequencing reads were mapped by BWA with a maximum of six mismatches and no gap [27]. Amplicons with the same tag were collected to generate a read cluster. Since each read cluster was originated from the same template, true mutations were called only if the mutations occurred in 90% of the reads withina a read cluster. Read clusters with a size below three reads were filtered out. Read clusters were further conflated into “error-free” reads. Relative fitness index (RF index) for individual point mutations was computed by:

(occurrence frequency in passaged library)/(occurrence frequency in plasmid library)

For all the downstream analysis, only point mutations covered with ⩾ 30 tag-conflated reads (“error-free” reads) in the plasmid library were included. This arbitrary cutoff filtered out mutants with low statistical confidence.

---

**Supplemental Figure 1.**
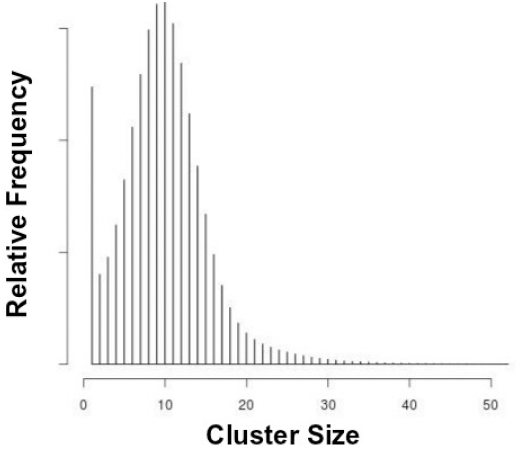
Distribution of conflated cluster size. Reads from the same amplicon with the same tag was defined as a cluster. The counts (number of reads) for all clusters are displayed as a histogram. Individual molecules, each carrying a unique tag, have an average copy number of ∼10 in the sequencing data, thus validating the sequencing library preparation design.

**Supplemental Figure 2.**
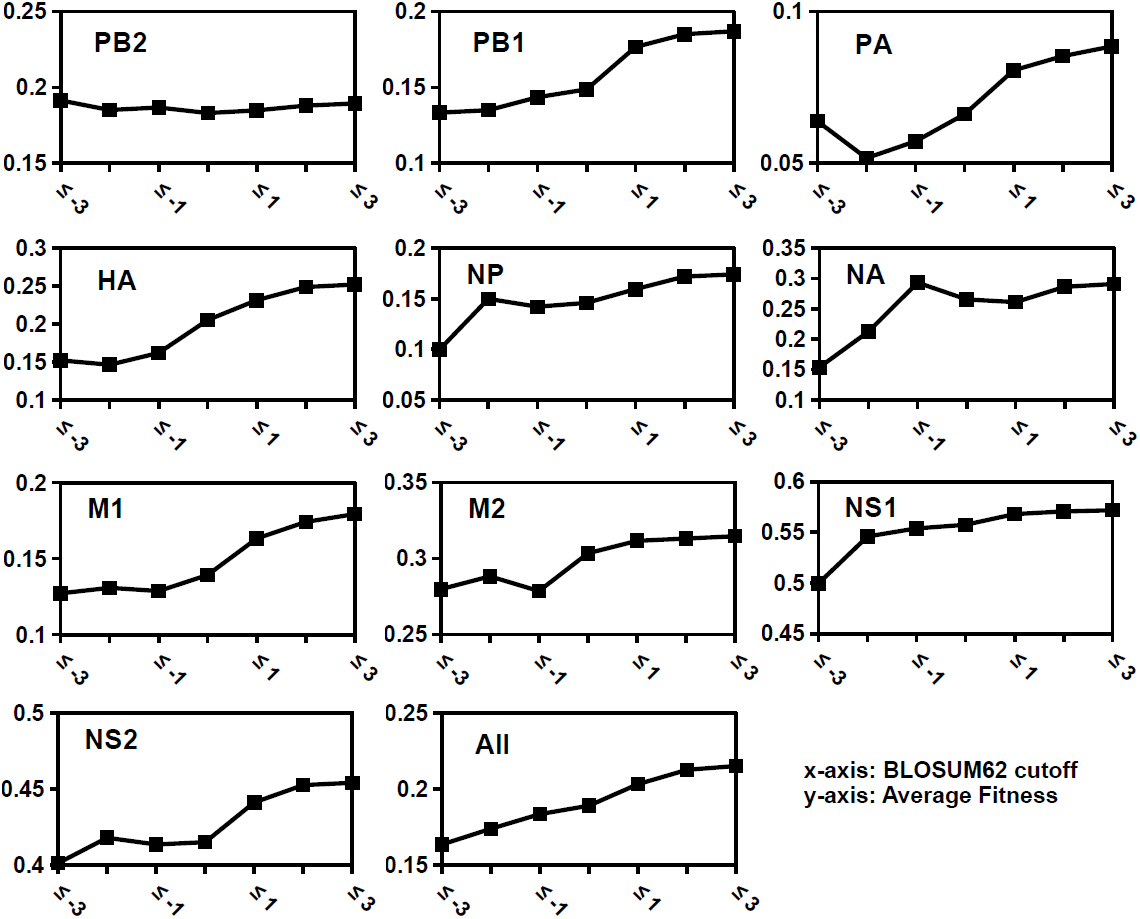
Comparison with BLOSUM62-based amino acid conservation. RF index of missense mutations from different segments were extracted and compared to amino acid conservation. The degree of amino acid conservation was quantified by the BLO-SUM62 matrix, a substitution matrix based on an implicit model of evolution. The x-axis represents the different cutoffs for BLOSUM62 values. The average RF index value for missense mutations that satisfied the cutoff was plotted against different BLOSUM62 cutoff values. The positive correlation between the RF index and the degree of amino acid conservation of missense mutations indicates that our fitness data shows consistency with the evolutionary trend for missense mutations.

**Supplemental Figure 3.**
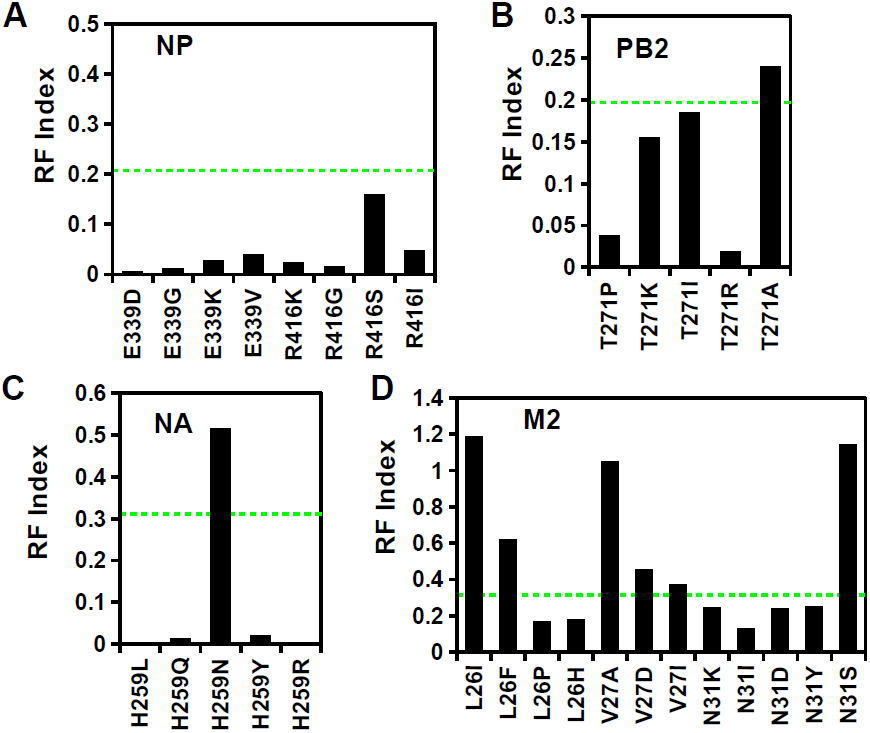
The RF index of substitutions at different functional sites. (A) E339 and R416 on the NP protein form a salt bridge at the homodimer interface, which is essential for viral replication [11]. This suggests that it is a feasible drug target. Several small molecules have been identified to target this interface and inhibit viral replication. (B) T271A has been identified as the replication enhancement substitution on PB2. T271A virus showed enhanced growth as compared to the WT strain in mammalian cells *in vitro* [12]. (C) NA 259Y (N1 naming: H274Y), a known oseltamivir drug resistance substitution, was shown to present a strongly attenuated phenotype in WSN [13]. In contrast, H259N (N1 naming: H274N), did not impose a deleterious effect in our fitness profiling data. This substitution is hypothesized to reduce influenza zanamivir sensitivity. Our results suggest further characterization of this substitution is warranted. (D) L26I, L26F, V27A and S31N on M2, the amantadine/rimantadine resistance substitutions [14, 15], were shown to impose little effect on viral replication. Our data is consistent with the observation that resistance substitutions emerged rapidly during amantadine/rimantadine drug treatment [5]. Green dotted line represents the average RF index for missense mutation at the indicated segment. Overall, the fitness data was consistent with the phenotypes of functional mutants reported in the literature.

**Supplemental Figure 4.**
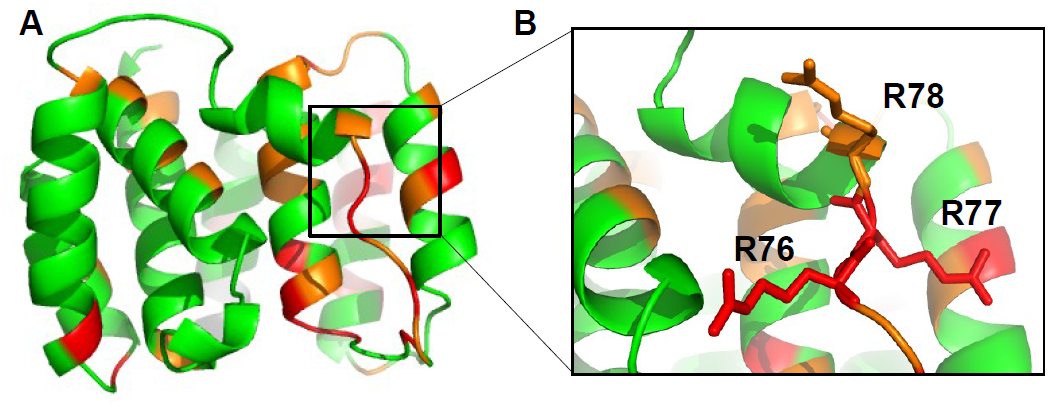
Structural analysis of M1. (A) The RF index of the least destructive missense mutations for individual amino acids on the M1 segment were projected on the protein structure (PDB: 1EA3) to identify indispensable regions [30]. The RF index was color coded: RF index < 0.1, red; 0.1 ⩽ RF index < 0.2, orange. (B) The critical residues _76_RRR_78_ were displayed in stick format as an inset. It has been suggested that this basic amino acid stretch is important for virus assembly and/or budding [16]. Virus substitutions at these positions show an attenuated phenotype. Our data is consistent with the previous observation. The non-structural region at the C-terminal end of _76_RRR_78_ is also indispensable in our profiling data. This suggests that entire the non-structural region containing the _76_RRR_78_ basic stretch is functionally important. One possibility for functional importance is that it provides an interface for a protein-protein interaction.

**Supplemental Figure 5.**
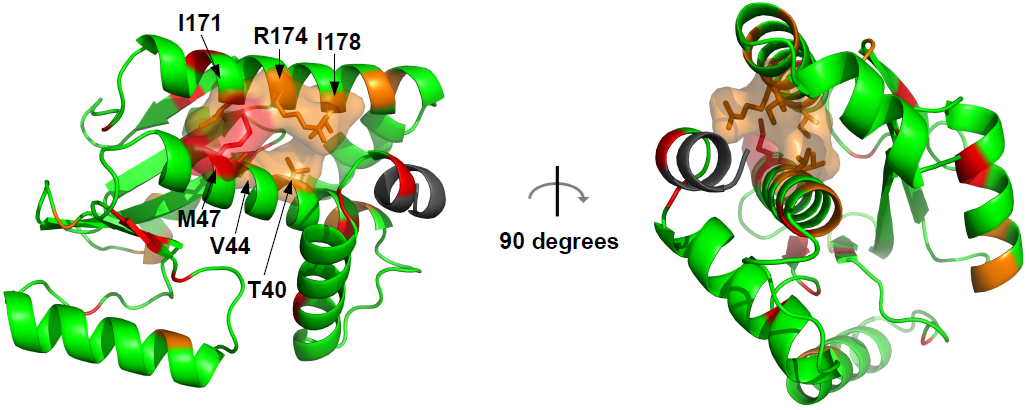
Structural analysis of the PA endonuclease domain. The RF index of the least destructive missense mutations in the profiling data for individual amino acids on the PA segment are projected on the PA endonuclease crystal structure (PDB: 4E5G). Most deleterious 10%, red; 10% to 20%, orange; Others, green. A critical helix-helix interface, which consists of T40, V44, M47, I171, R174 and I178, is highlighted. It demonstrates the power of qHRG in identifying residues that are not continuous in the primary sequence.

**Supplemental Figure 6.**
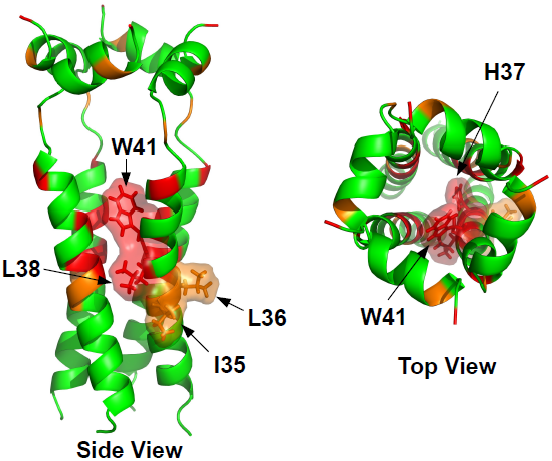
Structural analysis of the M2 ion channel. The RF index of the least destructive missense mutations in the profiling data for individual amino acids on the M2 protein are projected on the M2 ion channel crystal structure (PDB: 2RLF) [31]. Most deleterious 10%, red; 10% to 20%, orange; Others, green. An indispensable region on the transmembrane helix is highlighted. Our data captured the essential amino acids W41 and H37, which are critical for M2 ion channel activation [31]. We also identified several adjacent hydrophobic residues, I35, L36, and L38 as critical residues, which can be attributed to their contact with the hydrophobic membrane.

